# Separation and Characterization of Endogenous Nucleosomes by Native Capillary Zone Electrophoresis – Top-Down Mass Spectrometry (nCZE-TDMS)

**DOI:** 10.1101/2020.11.25.398925

**Authors:** Kevin Jooß, Luis F. Schachner, Rachel Watson, Zachary B. Gillespie, Sarah A. Howard, Marcus A. Cheek, Matthew J. Meiners, Jonathan D. Licht, Michael-Christopher Keogh, Neil L. Kelleher

**Author notes:** **Correspondence:** Prof. Dr. Neil Kelleher, 2070 Campus Dr, Evanston, Il 60208, USA.

## Abstract

We report a novel platform (native capillary zone electrophoresis – top-down mass spectrometry; nCZE-TDMS) for the separation and characterization of whole nucleosomes, their histone subunits, and PTMs. As the repeating unit of chromatin, mononucleosomes (Nucs) are a ~200 kDa complex of DNA and histone proteins involved in the regulation of key cellular processes central to human health and disease. Unraveling the covalent modification landscape of histones and their defined stoichiometries within Nucs helps to explain epigenetic regulatory mechanisms. In nCZE-TDMS, online Nuc separation is followed by a three-tier tandem MS approach that measures the intact mass of Nucs, ejects and detects the constituent histones, and fragments to sequence the histone. The new platform was optimized with synthetic Nucs to reduce both sample requirements and cost significantly compared to direct infusion. Limits of detection were in the low attomole range, with linearity over ~three orders of magnitude. The nCZE-TDMS platform was applied to endogenous Nucs from two cell lines distinguished by overexpression or knockout of histone methyltransferase NSD2/MMSET, where analysis of constituent histones revealed changes in histone abundances over the course of the CZE separation. We are confident the nCZE-TDMS platform will help advance nucleosome-level research in the fields of chromatin and epigenetics.

## 1. Introduction

Increasing evidence indicates that disruption in the cellular epigenetic machinery may be a key initiator in several types of cancer and other disorders.^1^ The mononucleosome (Nuc), the smallest repeating unit of chromatin, resides at the center of epigenetic regulation. It comprises ~147 bp of DNA wrapped around a protein octamer with two copies of each core histone: H2A, H2B, H3, and H4 (**Figure S1a**).^2^ Nucleosome structure (and presumably function) can be altered by covalent modification of the DNA or histone proteins (latter termed post-translational modifications; PTMs), or by replacement of the canonical histones with variants, which have diverse biophysical properties.^3, 4^ Nuc modification dynamics can regulate key cellular processes central to human health and disease (e.g. transcription, DNA repair, and DNA replication).^5^ For this reason, the notion of a ‘histone code’ has gained traction over the last 20 years, given the discovery of effector proteins that simultaneously recognize and bind one or more PTMs on nucleosome assemblies.^6^ Unraveling the full landscape of modifications on whole Nucs is thus key to elucidate the mechanisms that exert epigenetic control over a genomic location.

Chromatin and epigenetics researchers generally use histone peptides as Nuc proxies when studying PTM writers, readers and erasers, obscuring the combinatorial variations that co-occur at the Nuc-level and mediate genome transactions.^7, 8^ The high cost of novel PTM-defined Nuc reagents and lack of a direct compositional readout have slowed Nuc-level biology.^9, 10^ To address this technology gap, we recently described “Nuc-MS”, a novel method based on top-down mass spectrometry (TDMS) operated in native mode, that can decode the protein composition of whole nucleosomes from synthetic and endogenous sources.^11^ A major feature of TDMS is the isolation and fragmentation of individual histones, such that PTMs and variant substitutions can be directly assigned as a single nucleosome proteoform.^12^ The top-down approach has been extended to whole complexes by our group and others^13–17^ and provides a higher-level view of the modification landscape compared to the inference required in bottom-up approaches.^18^

In recent years, native mass spectrometry (nMS) has greatly contributed to structural biology research.^19, 20^ Here, volatile aqueous buffer systems (e.g. ammonium acetate) are employed to preserve near-native features of proteins and their complexes during electrospray ionization.^21, 22^ The approach allows to evaluate the composition of biomolecular structures leading to a better understanding of their biological function and significance.^15, 22^ In particular, coupling native ionization with TDMS provides insights to subunit stoichiometry, stability, topology, dynamics, and the affinities of protein complexes.^23^ The lower extent of protonation during native electrospray ionization (ESI) gives access to a wider range of the MS instrument’s *m/z* region than under denatured ESI, aiding the resolution of different protein signals.^24^ However, ion suppression and signal superposition cannot be completely avoided for complex samples. Thus, upfront separation under native conditions is beneficial and highly desired.

Different analytical chromatographic^25–28^ and electromigrative^29–31^ tools theoretically allow native separation. Generally, there are three major requirements to achieve efficient native separation prior to MS detection: (i) the mobile phase or background electrolyte (BGE) must maintain the nature of the non-denatured protein or complex; (ii) the composition of the mobile phase/BGE must be compatible with electrospray ionization; and (iii) the separation performance and resolution needs to be sufficient. Unfortunately, many current techniques suffer in at least one of these aspects. In contrast, capillary zone electrophoresis (CZE) provides high-resolution even under native conditions.^23^ The separation mechanism of CZE is based on the electrophoretic mobilities of ions in the liquid phase, which are dependent on the charge-to-size ratio of the analytes.^32^ CZE has been shown to be well-suited for the separation of intact proteins with only small structural differences such as deamidation events.^33^ Another important aspect of CZE is the low sample requirement, typically a few nanoliters per injection.^34^ Several CZE-MS interfaces have been recently developed, including sheath liquid, nanoflow sheath liquid and sheathless porous-tip.^35^ However, native (n)CZE-TDMS for protein analysis remains a largely unexplored area, with few studies to date.^31, 36, 37^

The presence of PTMs and histone variants is generally asserted through immunoblots, affinity-based enrichment (e.g. ChIP-seq), and digestion-based proteomics.^38^ Here, we report the first native-mode MS platform for the separation and characterization of intact Nucs, their histone subunits and PTM profiles in a single experiment. The nCZE-TDMS method was initially optimized using synthetic Nucs to enable MS^1^, MS^2^ and pseudo-MS^3^ data collection (**Figure S1b**), and later expanded for the separation of endogenous nucleosomes (endoNucs) using methyltransferase-modulation cell lines as a model system. An important feature of the platform is that Nuc samples in complex buffers were directly injected into the CZE system without any sample preparation, only consuming minor amounts of sample material (<$0,01 per injection). As a result, sample requirements - and thus cost - are significantly reduced compared to direct infusion, enabling the online separation of Nuc sub-populations based on their charge-to-size ratios. This new data type, coupled with Nuc separation by CZE, provides a wide-lens, semi-quantitative view of a cell’s epigenetic landscape.

## 2. Materials and methods

### 2.1 Materials

Water (Optima^®^ LC-MS grade), acetic acid (HAc) glacial (Optima LC-MS grade), and hydrochloric acid (HCl) (technical grade) were from Thermo Fisher Scientific (Chicago, IL, USA). 7.5 M ammonium acetate (AmAc) stock solution (molecular biology grade) was from Sigma Aldrich (St. Louis, MO, USA). Molecular weight cut-off (MWCO) spin filters were from Thermo Fisher Scientific. All solutions used in the CESI 8000 Plus system were passed through 0.2 μm pore size Nalgene™ Rapid-Flow™ sterile disposable filters (Thermo Fisher Scientific).

### 2.2 Synthetic nucleosome samples

All semi-synthetic nucleosomes were from EpiCypher (Durham, NC, USA): unmodified recombinant (r)Nuc (16-0006); H3K27me3 (16-0317), H2BK120ub (16-0370) and H3K4,9,14,18ac (16-0336) designer (d)Nucs; and H3.3G34V (16-0347) oncoNuc. For CZE experiments, Nuc samples were kept in their original storage buffer (10 mM Tris-HCL, 1 mM EDTA, 25 mM NaCl, 2 mM DTT and 20% glycerol) and only mixed and/or diluted in BGE as needed.

For direct infusion experiments via the CESI 8000 Plus device, Nuc samples were buffer exchanged using Amicon Ultra-0.5-mL centrifugal filters (30 kDa MWCO). In brief, the filter device was equilibrated with 500 μL of 50 mM AmAc and spun for 5 min at 12,000 ×g. The Nuc sample was then loaded up to a total volume of 500 μL, and spun for 5 min at 12,000 ×g or until concentrated to ~50 μL. Eight consecutive washing steps were performed by adding AmAc (40 mM) to 500 μL and spinning at 12,000 ×g for 5 min. The remaining sample volume was transferred to CESI NanoVials for analysis.

### 2.3 Endogenous nucleosome samples

TKO and NTKO NSD2 low and high expressing cells were prepared as described,^39^ with endogenous Nucs extracted and prepared for MS analysis as previously.^11^ The only deviating step was a final buffer exchange to 40 mM AmAc for nCZE-TDMS analysis, resulting in a concentration of about 10 mg/mL of endoNuc material.

### 2.4 Capillary Electrophoresis

A CESI-8000 Plus instrument from SCIEX (Redwood City, CA, USA) was used, and separation was performed using commercial Neutral OptiMS™ Capillary Cartridges (30 μm inner diameter, 90 cm length) containing an integrated sheathless etched porous nanospray tip. The inner wall of the separation capillaries is coated with a binary layer: (i) hydrophobic coating and (ii) hydrophobic polyacrylamide surface. In this way, protein adsorption is prevented, and electro-osmotic flow is largely suppressed.

For initial conditioning, the capillary was rinsed (100 psi) with 0.1 M HCl (5 min), BGE (10 min) and water (30 min); the conductive line (CL) rinsed with water (5 min). Subsequently, both separation and conductive line were rinsed (100 psi) with BGE (5 and 3 min, respectively) followed by applying high voltage of+15 kV and 5 psi supporting flow for 60 min. At the end of this step, the voltage was ramped down over 5 min. Each morning, the capillary was rinsed with fresh BGE and high voltage (+15 kV, 5 psi) was applied for 30 min. For long term storage, capillaries were rinsed with water and kept at 4 °C.

For Nuc analysis, the capillary was first rinsed with 0.1 M HCl (100 psi, 3 min), followed by BGE (5 min). The CL was filled with 3% HAc (100 psi, 3 min). Hydrodynamic injection was performed at 2.5 psi for 30 sec, corresponding to ~6 – 38 nL (0.9 – 6.0% of the total capillary volume), followed by a water dipping step and the injection of a small plug of BGE (2.5 psi, 10 sec). Three different CE methods were developed during this work: (i) standard; (ii) high-resolution; and (iii) high-throughput. A detailed description of each individual method is in **Table S1**.

### 2.5 Mass Spectrometer

The CESI-8000 Plus instrument was hyphenated with a custom Thermo Fisher Q Exactive Orbitrap HF MS with Extended Mass Range^14^ (QE-EMR) and a commercial Thermo Fisher Q Exactive Orbitrap MS with Ultra High Mass Range (UHMR). Important parameters of the applied tune files for MS^1^, MS^2^, and pseudo-MS^3^ experiments for both instruments are in **Table S2**. A Nanospray Flex™ Ion Source was changed using an OptiMS Thermo MS Adapter from SCIEX for hyphenation of the CESI and orbitrap (OT) instruments. The sprayer tip was positioned 2.5 mm in front of the OT orifice and an ESI voltage between +1.6 and +1.9 kV was applied during separation. The inlet capillary of the MS instruments was heated to 330 °C (QE-EMR) and 300 °C (UHMR), respectively.

### 2.6 Data Analysis

Spectra were analyzed manually using Thermo Xcalibur 4.0 Qual Browser (Thermo Fisher Scientific, Inc.). Figures were created using Adobe Illustrator CC 2015.3. Signal-to-Noise ratios (SNR) were calculated as follows: SNR = (NL – B) / (N – B), where NL is the signal intensity, B is the baseline intensity, and N is the noise intensity. The sum of all observed charge states for each parameter (NL, N and B) was used for SNR calculations. In this way, an average SNR for the protein charge state distribution was determined. Least-square regression was performed, and the residual errors were weighted 1/x, where x is the respective concentration level. Limit of Detection (LOD, SNR = 3) was estimated by extrapolation of the lowest measured concentration level. Deconvolution of histone data (MS^2^) was performed using Unidec 3.2.0.^40^ Parameters: Data processing: Range 500 – 2500 Th, Bin every: 0; Charge Range: 5 – 15, Mass Range: 10 – 20k Da, Sample mass every (Da): 0.05. Peak area values were calculated by integrating the assigned mass range of proteoforms in the deconvoluted mass spectrum. Histone proteoforms were assigned by intact mass and isotopic fit (in-house data base). Statistical significance was evaluated using two-sided, two-sample t-tests and the resulting p-values compared to Bonferroni corrected α-values based on α = 0.05%. Mass lists of peptide fragments were created using Xtract (Thermo Fisher Scientific). TDValidator^41^ (max ppm tolerance: 16 ppm; sub ppm tolerance: 7.5 ppm; cluster tolerance: 0.35; charge range: 1-15; minimum score: 0.3; S/N cutoff: 3; Mercury7 Limit: 0.0001; minimum size: 2) was used to assign recorded fragment ions to the primary sequence of the histone subunits. Electrophoretic resolution was calculated based on the full width half maximum of the respective CZE peaks.

## 3. Results and Discussion

We first evaluated the performance of the nCZE-TDMS platform with a recombinant Nuc containing the histone H3 trimethylated at lysine 27 (H3K27me3, c = 1 μM, **Figure 1a-c**). Importantly, the sample was injected directly in the manufacturers storage and shipping buffer (10 mM Tris-HCL, 1 mM EDTA, 25 mM NaCl, 2 mM DTT, 20% glycerol: EpiCypher) which was designed to minimize Nuc loss and degradation. The data show that both salts and other buffer components are clearly separated from the Nuc signal in the total ion electropherogram (TIE, **Figure 1b**), obviating the need for additional sample purification such as solvent exchange to AmAc solutions.^42^ A general scheme of the separation system is depicted in **Figure 1d**. The injection volume was subsequently optimized (**Figure S2**), and 2.5 psi for 30 s (~11 nL) selected for further experiments. Comprised of both basic histones (pI >10) and acidic DNA strand (pI ≈5)^43^, the effective pI and mobility of Nucs are hard to predict. Therefore, the separation voltage was varied between +12 and +18 kV while keeping the remaining method parameters constant to evaluate the migration behavior of Nucs under native conditions (**Figure S3**). Higher voltages resulted in an increase in migration time, indicating the net charge of the Nuc complexes is negative under the given conditions. Consequently, in the applied positive polarity mode they would counter-migrate towards the CESI inlet, but the supplemental pressure is sufficient to drive the Nucs towards the electrospray source.

**Figure 1:**
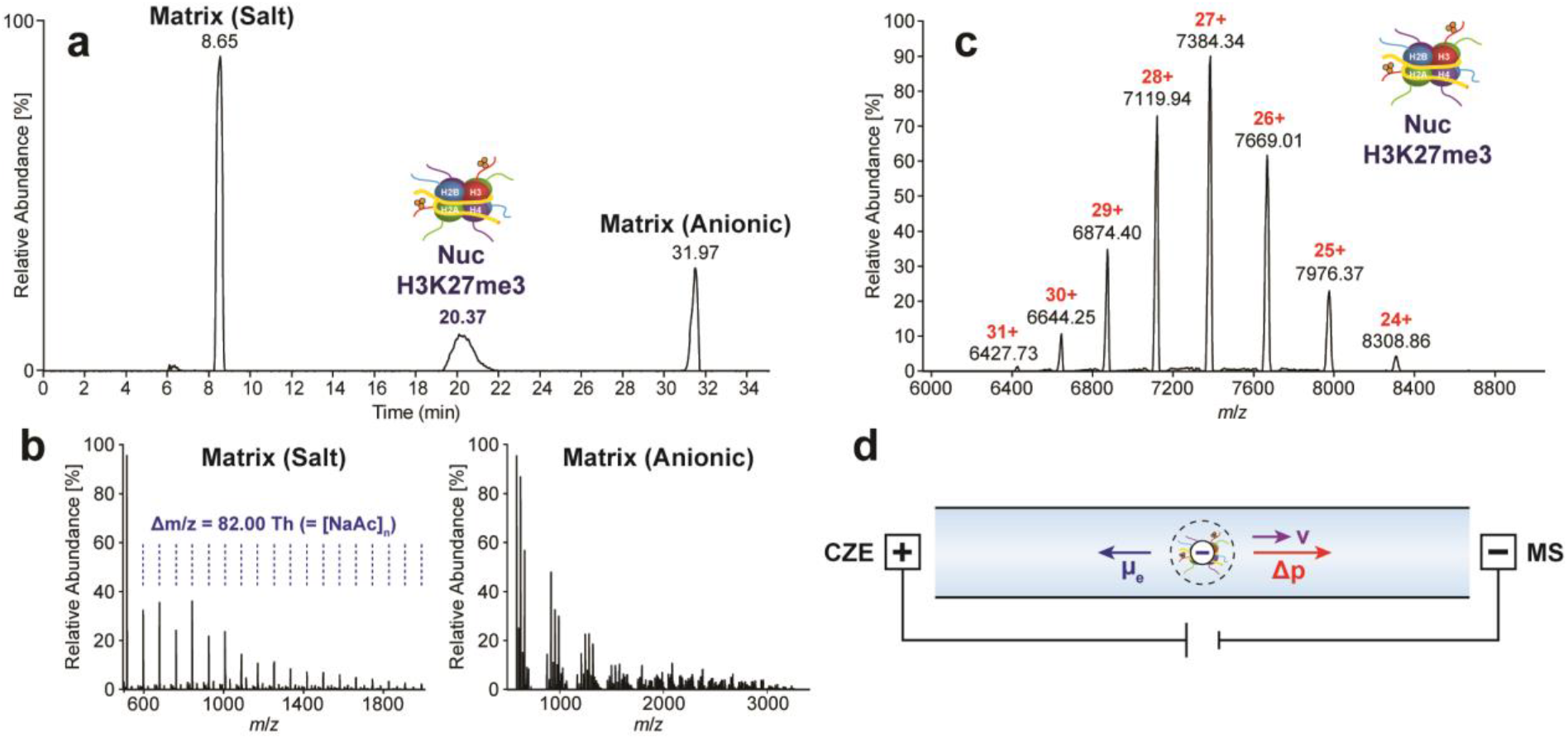
Native CZE-TDMS analysis of Nucs. (a) TIE of Nuc containing H3K27me3 (c = 1 μM) showing the separation of analyte signal and matrix. (b) Mass spectra of high-intensity matrix compounds including sodium acetate clusters. (c) Mass spectrum of intact H3K27me3 nucleosome. (d) Scheme of Nuc migration during analysis. Nucs are negatively charged in AmAc based BGEs (pH ≈6.8)

We next tested histone ejection from the intact Nucs using this platform. An example MS^2^ spectrum of H3K27me3 Nucs after ion-source ejection (c = 1 μM, **Figure 2a)** demonstrates detection of the four histone subunits H4, H2B, H2A, and H3K27me3 (**Figure 2b**). These ejected histones were subsequently isolated and fragmented by higher-energy collisional dissociation (HCD)^14^ and identified by TDValidator (e.g. H2A: **Figure 2c-d**). Of note, the quality of fragmentation from online CZE-TDMS was comparable to native direct infusion analyzing the same sample (**Figure S4**). Another example showing H3K27me3 fragmentation can be found in **Figure S5**. The native TD approach assigns ejected histones to their related precursor Nuc. Thus, this procedure constitutes an important gain of information relative to denaturing or proteolysis-dependent histone analyses.^44^

**Figure 2:**
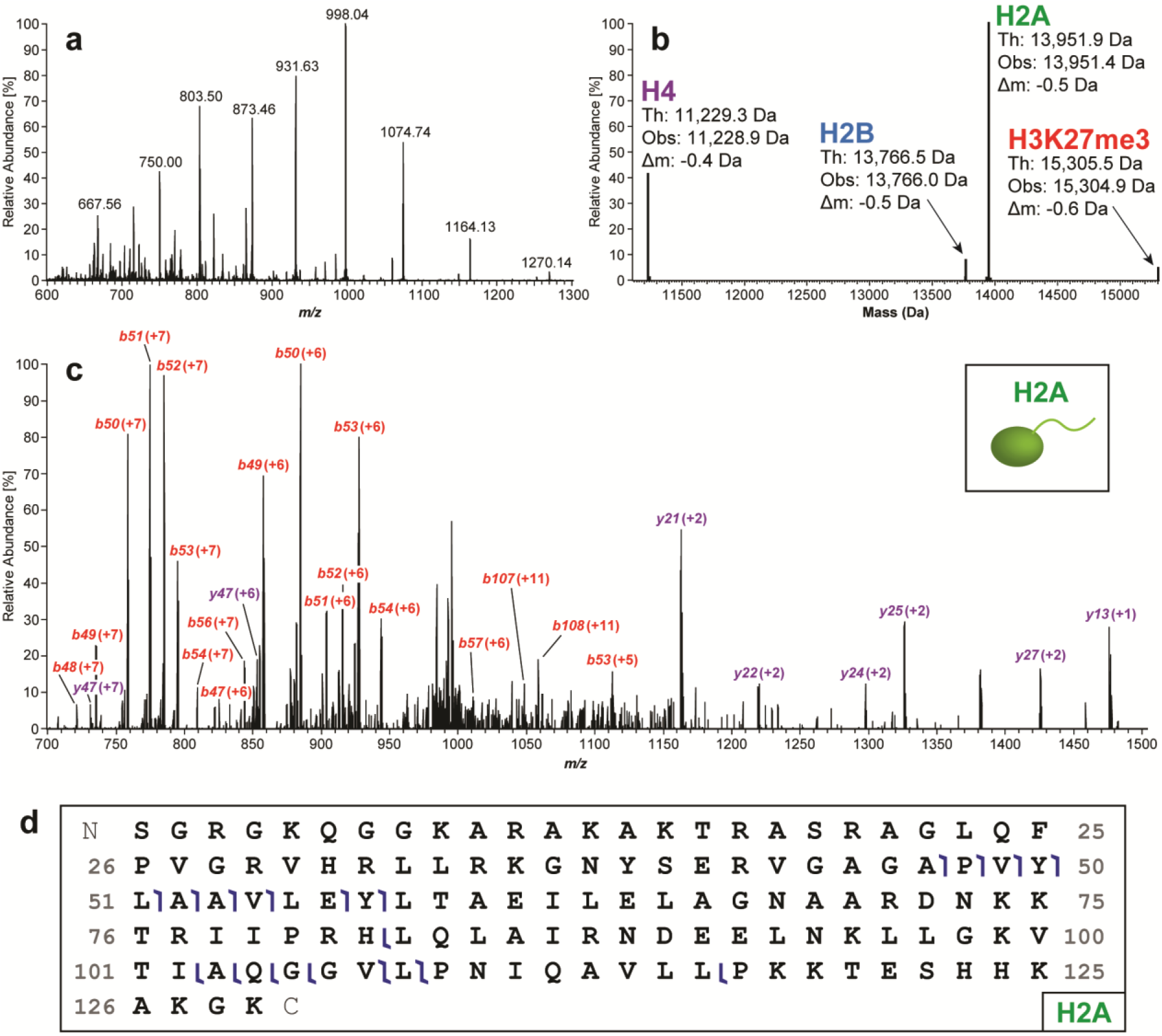
Native CZE-TDMS analysis of H3K27me3 (c = 1 μM). (a) Mass spectrum of histones ejected from intact Nucs. (b) Deconvoluted mass spectrum of ejected histones including H4, H2B, H2A, and H3K27me3. (c) After histone ejection from the Nuc using in-source dissociation, individual histones were isolated and fragmented by HCD. An example fragment mass spectrum of H2A is depicted. (d) Graphical fragment map of peptides observed from H2A.

The nCZE-TDMS platform shows high intra- and inter-day reproducibility in migration time, peak area, and intensity using synthetic Nucs, indicating its potential to support the industrial production / analysis of such reagents (**Table S3**). Moreover, we tested LOD and quantitation at the MS^1^ level, finding that the linear range of detection is between 10 nM and 500 nM (R^2^ = 0.996), with a SNR of 11.3 ±1.0 (10 nM Nuc sample; = 110 amol), resulting in an estimated LOD of 2.7 nM (≙ 30 amol) (**Figure S6a-c**). In a follow-up experiment we showed adequate MS^2^ data for ejected histones from unmodified synthetic Nucs at a concentration of 62.5 nM (≙ 0.71 fmol, **Figure S6d**).

We next sought to investigate the Nuc separation capabilities of the CZE-TDMS platform. For proof-of-concept, we mixed tetra-acetylated (H3K4,9,14,18a) and ubiquitinated (H2BK120ub) Nucs prepared (500 nM each) as models of highly divergent Nuc species. Tetra-acetylation / charge neutralization of eight lysine residues (i.e. both histone H3 tails) will decrease the overall net charge of the Nuc complex. The mass shift introduced by two ubiquitins is rather large (2 × 8.5 kDa), but it is challenging to predict its influence on the overall net charge of the Nuc complex (the ~6.8 pI of ubiquitin is close to the pH of the developed CZE method). The mixture was analyzed using the “standard” CZE settings (**Figure 3a**), achieving partial separation (R = 0.366, n = 2) with H2BK120ub detected first. Under different parameters (evaluated considering separation performance; **Discussion S1**) migration times and peak widths increase noticeably when decreasing the supplemental pressure from 5.0 psi to 2.2 psi, electrophoretic resolution is significantly improved from R = 0.37 to 1.09 (n = 2 each, **Figure 3b**), which becomes even more evident by comparing the mass spectra in **Figure 3c-f.** The SNR also improved by ~three-fold, likely due to reduced ion suppression from overlapping peaks and a higher number of scans averaged.

**Figure 3:**
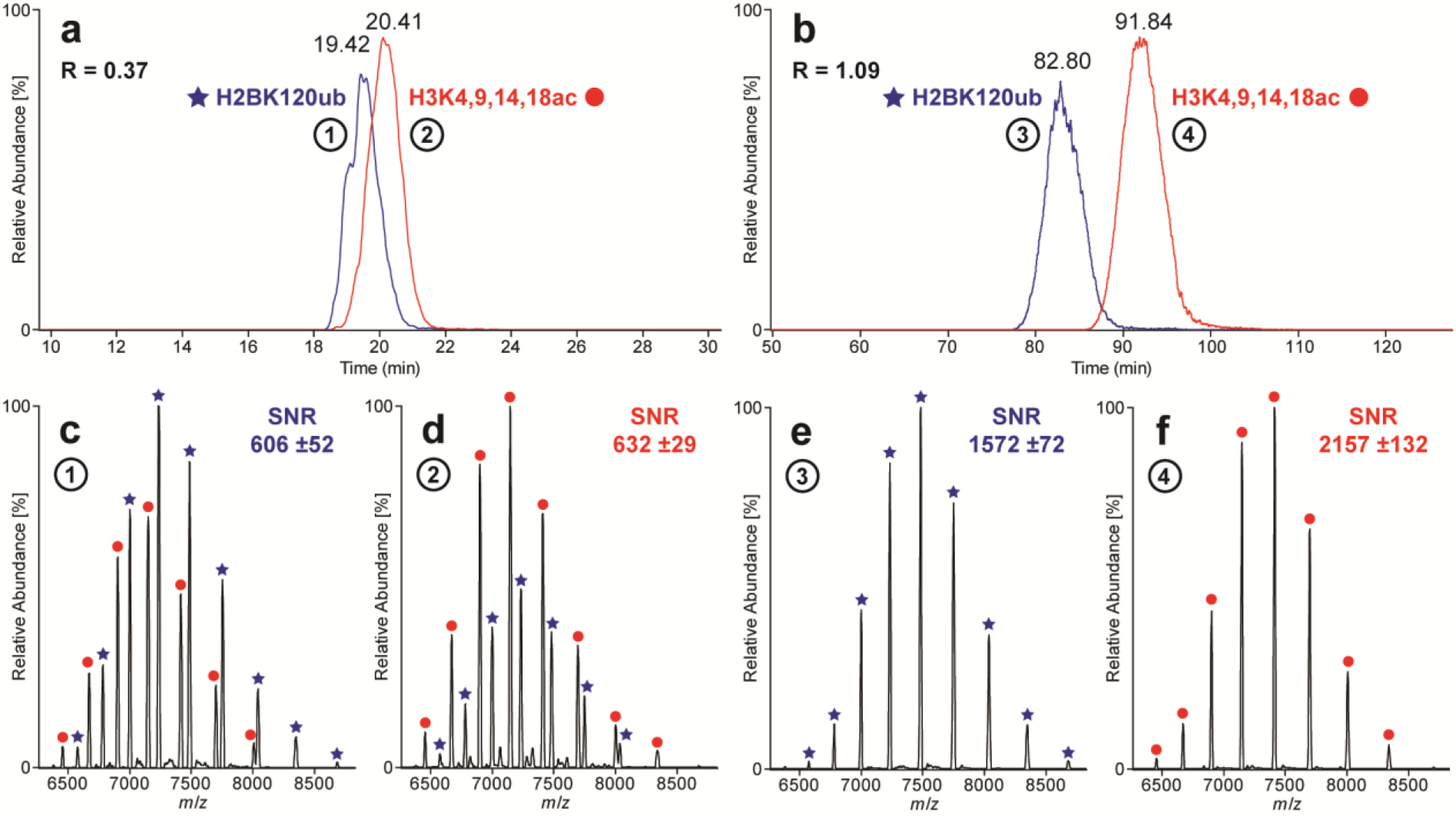
Comparison of “standard” and “high-resolution” CZE methods. Extracted Ion Electropherograms (EIEs, seven highest charge states each) of sample containing H2BK120ub and H3K4,9,14,18ac analyzed with (a) standard and (b) high-resolution CZE method. Mass spectra (c-f) generated by averaging the EIE peaks at full width at half maximum (FWHM). Partial separation (R = 0.366, n = 2) obtained using standard method and close to baseline separation (R = 1.082, n = 3) achieved for high-resolution method.

In short, the final “high-resolution” CZE method is based on 40 mM AmAc (pH = 6.8) as BGE and a separation voltage of +18 kV with 2.2 psi of supplemental pressure, resulting in a total run time of about 135 min. With potential for widespread deployment of the nCZE-TDMS platform, we also developed a “high-throughput” CZE method with a total run time of ~20 min tailored for the quality control environment during Nuc manufacturing (see **Discussion S2**).

As a next step, the native CZE-TDMS platform was applied for the analysis of endogenous nucleosomes (endoNucs) derived (see **Methods**) from an isogenic knockout and overexpression system (respectively TKO and NTKO cell lines^45, 46^) of NSD2/MMSET, a histone methyltransferase implicated in t(4;14)+ multiple myeloma pathogenesis. In TKO cell lines, the rearranged IGH-NSD2 allele is inactivated by homologous recombination, leaving one wild-type copy of the NSD2, which yields low levels of H3K36me2 (NSD2^+/-^; H3K36me2^lo^). In NTKO cells, the wild-type NSD2 allele is inactivated leaving the IGH-NSD2 allele and high levels of H3K36me2 (IGH-NSD2; H3K36me2^hi^).^45^, ^46^

The complexity of endoNucs cannot be successfully resolved by traditional direct infusion nMS, which yields a low-abundant and indistinct protein signal (data not shown). In the nCZE-TDMS system, endoNucs showed increased migration time relative to synthetic Nucs. This indicates a higher electrophoretic mobility, so the supplemental pressure was increased from 2.2 to 3.0 psi (**Figure 4a** & **Figure S7**). The endoNucs migrated over a large time window (~70 to 80 min), which was divided into five-minute subsections and individually deconvoluted to search for distinct populations.

**Figure 4:**
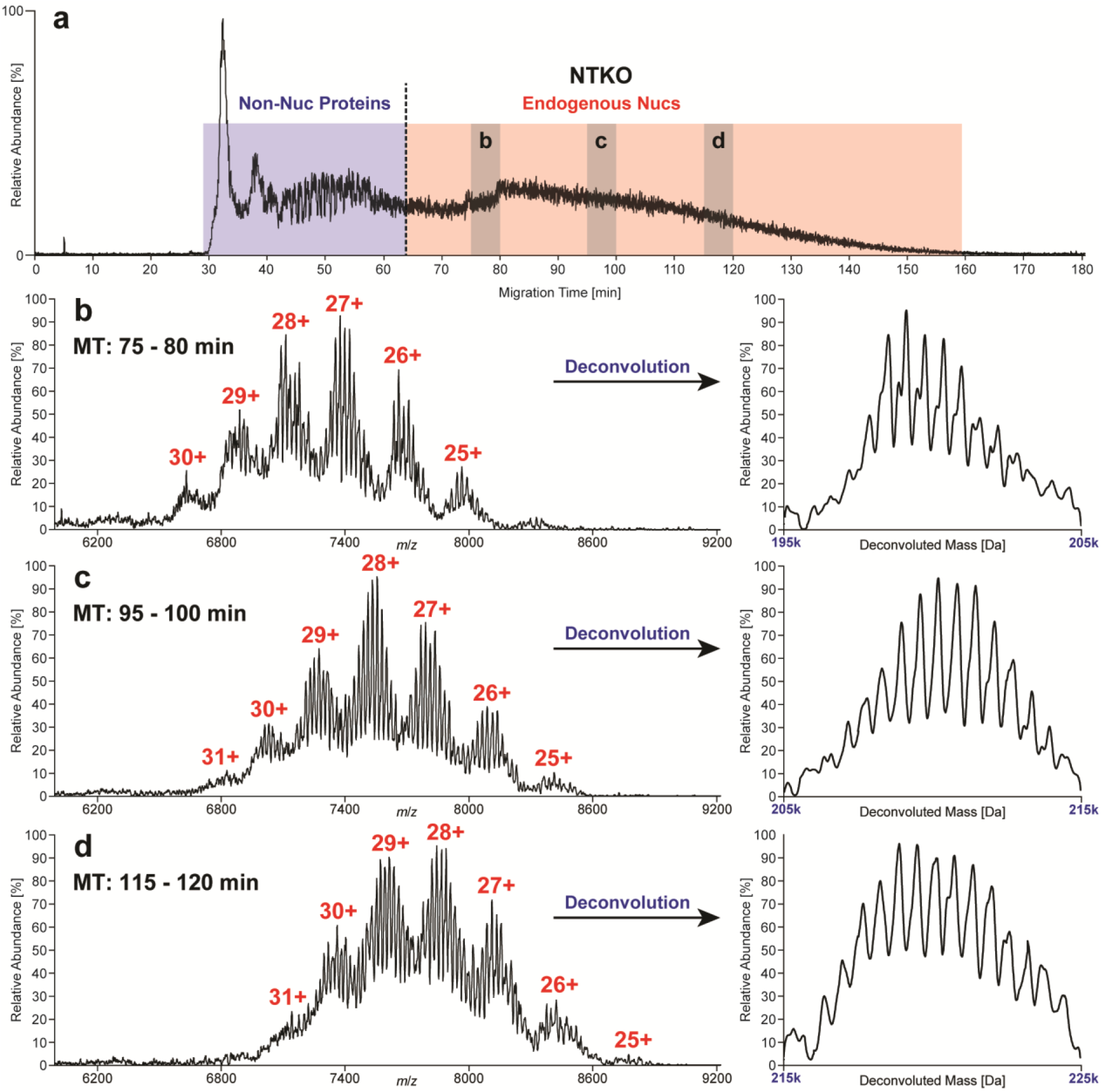
Separation of endoNucs (~10 mg/mL) derived from NTKO (IGH-NSD2; H3K36me2^hi^) cells. TIE (a) divided into two major regions: (i) non-Nuc proteins and (ii) endogenous Nucs. Raw and deconvoluted mass spectra (b-d) of three exemplary sections (75 - 80 min, 95 - 100 min, 115 - 120 min), where several “nucleoforms” predominated in each. In general, the intact mass of nucleoforms increased with migration time. The average mass shift of 620.2 ±18.5 Da between neighboring peaks, corresponds to a base pair difference in the length of the associated DNA (Δm_GC_ = 618.4 Da, Δm_AT_ = 617.4 Da).

The raw and deconvoluted spectra in **Figure 4b-d** reflect a variety of distinct peaks that increase in mass (192,332 to 228,854 Da) over the length of the CZE run. The mass differences between the detected peaks reveals up to 60 “nucleoforms” that differ by a base pair (NTKO: 620.2 ±18.5 Da; TKO: 617.5 ±22.4 Da; [Δm_GC_ = 618.4 Da, Δm_AT_ = 617.4 Da]); likely reflecting processive micrococcal nuclease (MNase) digestion products. In addition, a series of nonbaseline resolved satellite peaks were observed with a mass shift of ~300 Da, indicating single nucleotide differences (**Figure 4b**). These results represent the first report of base pair resolution and accurate mass measurement of intact Nucs from endogenous sources.

The same samples were analyzed at the histone level (MS^2^, n = 3 per sample), revealing significant proteoform-level differences between NTKO and TKO cells (**Figure 5a-d, Table S4**). Notably, we detected H3.2 as the predominant H3 subtype in these cells (proteoforms are labeled accordingly; **Figure S8**), although the presence of minor amounts of H3.1 cannot be excluded due to their closely related intact masses. The average methylation equivalent (ME) of H3.2 was decreased by ~11% in TKO cells, consistent with their reduced expression of NSD2 and lower levels of H3K36 methylation.^47^ Interestingly, the H3.2 4x ME proteoform is highly elevated in TKO, as compared to 6x and 7x ME in NTKO, suggesting NSD2 cross-talk with pre-existing PTMs. In this regard, H4 acetylation (most likely at K16)^11^ is 21% elevated in NTKO, suggesting increased areas of active transcription when NSD2 is overexpressed.^48^ Furthermore, the relative abundance of all H3.3 proteoforms is increased in NTKO cells, with 3x to 5x ME readily detected. Finally, the nCZE-TDMS platform directly quantifies H2A and H2B variants and their proteoforms (**Table S4**), revealing modification stoichiometry differences, with potential functional implications. Specifically, there is a higher degree of acetylation on H2A.1C and H2A.1B/E in NTKO versus TKO cells (17% and 48% increase respectively), consistent with the H4 observation. In terms of H2B, there is a lower abundance of variants H2B.1-K and H2B.1-C/E in NTKO relative to TKO cells, suggesting unexplored functional roles for H2B variants in epigenetic regulation.

**Figure 5:**
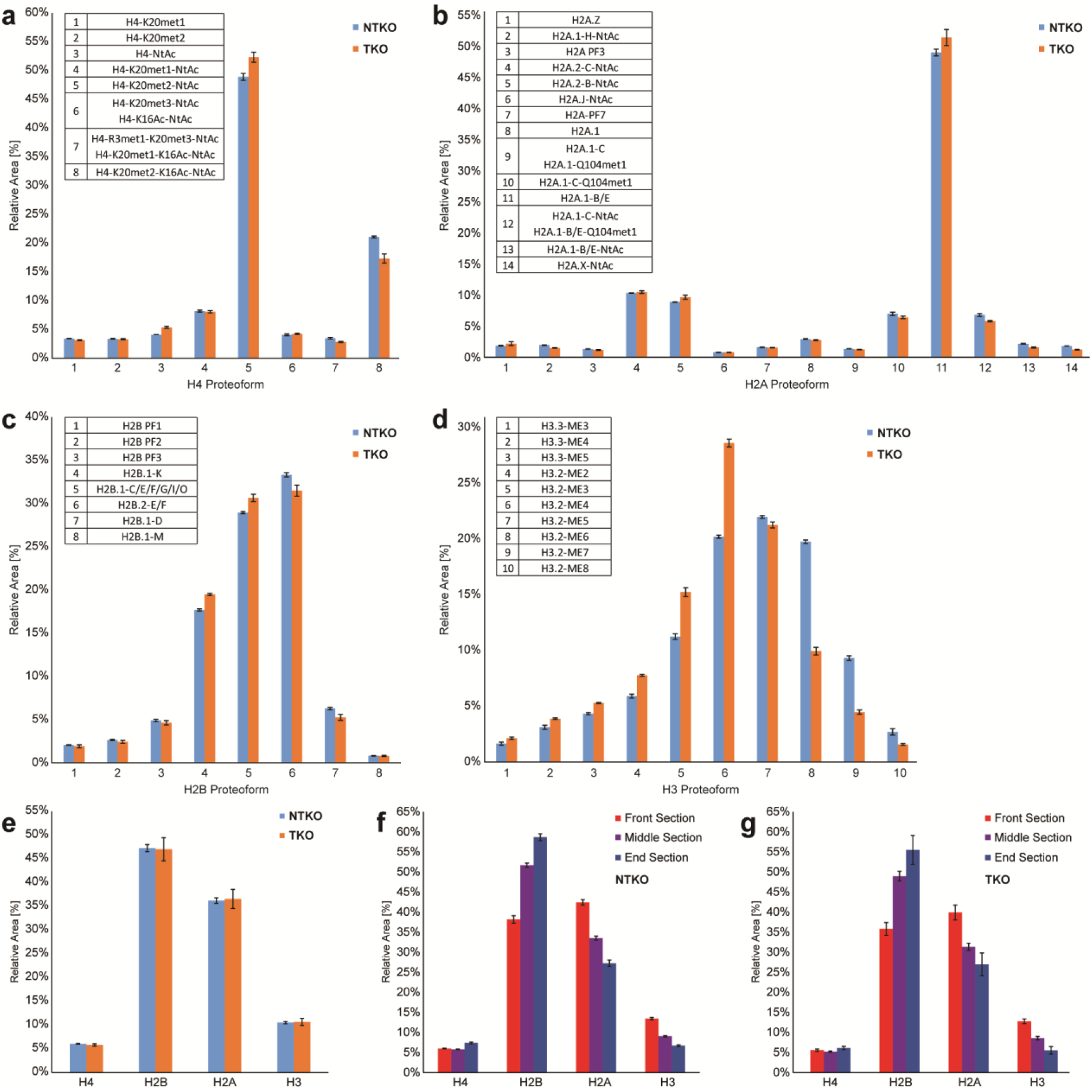
Relative quantification of NTKO/TKO histones using nCZE-TDMS (n = 3). (a-d) Histone proteoform level: Averaged mass spectra generated over the entire endoNuc migration range were used and the total peak area of H4 (a), H2A (b), H2B (c), and H3.1 (d) proteoforms was set to 100%, respectively. (e-g) Histone type level: (e) Comparison between NTKO (IGH-NSD2; H3K36me2^hi^) and TKO (NSD2^+/−^; H3K36me2^lo^) samples using spectra averaged over the entire endoNuc migration time. Comparison between front, middle and end section of the endoNuc migration time window for NTKO (f) and TKO (g). For this purpose, electropherograms were divided into three equally long sections.

Interestingly, time resolved data shows the relative ratio between histones change significantly over the course of the CZE separation (**Figure 5f-g**, **Table S5** & **S6**). In particular, we observed an increase in overall H2B, with a concomitant decrease in H2A and H3, as the Nuc particles increased in size at higher migration times. This is difficult to explain given our understanding of nucleosome structure and histone stoichiometry / exchange therein. However, inconsistent MNase digestion yields Nucs with varying DNA length (**Figure 4**), and it is possible that non-covalent interactions between histones and nucleotides in various nucleoforms differentially impacts efficiency of histone ejection (e.g. H2B is less tightly bound, explaining its elevated relative abundance with increasing DNA length).

At the proteoform level, some distinct changes over the course of the CZE separation were also observed (**Tables S7 & S8**). For example, an increase in H4-K20met2-NtAc (+14%) and H2A.1-B/E (+16%), but decrease in H4-R3met1-K20met3-NtAc / H4-K20met1-K16Ac-NtAc (−42%), H2A.1 (−34%), and H2A.1-C-Q104met1 (−30%) was detected at higher migration times for NTKO (and analogous changes in the proteoforms profiles for TKO).

## 4. Conclusion

In summary, we have developed the nCZE-TDMS platform to achieve attomole-level Nuc characterization with emphasis on either high resolution or throughput. In this proof-of-concept study, we demonstrate its potential to separate Nucs containing different proteoforms, which has not been shown by other techniques to date. With $0.01 of material consumed per 20 min run and minimal sample preparation, the system is a promising candidate for quality control of semi-synthetic Nucs. We challenged the platform with endoNucs from an isogenic methyltransferase knock-out / overexpression cell line system (NTKO/TKO), detecting changes in the H3 methylation profile consistent with NSD2 status, and proteoforms indicating potential cross-talk with other PTMs, including H4K16ac. Considering CZE separation performance, at the MS^1^ level we achieved base-pair resolution and detected ~60 distinct “nucleoforms”, associated with differential enzymatic cleavage of DNA by MNase. At the histone level (MS^2^ data) changes in proteoform abundances over the course of nCZE separation were observed for both NTKO and TKO, demonstrating the capabilities to resolve different Nuc sub-populations. We envision extending into the area of endogenous Nucs from cells and immuno-precipitated specimens where limiting amounts of starting material are available. In future studies, we aspire to simplify the complexity of nucleoforms by optimizing the digestion process to get a better overview of proteoforms changes. In conclusion, nCZE-TDMS has the potential to serve as an important tool in the field of epigenetics, native proteomics^17^ and in the quality control of complex synthetic biomolecules.

## Acknowledgements

This work was supported by the National Institute of General Medical Sciences P41 GM108569 for the National Resource for Translational and Developmental Proteomics at Northwestern University and NIH grants S10 OD025194 and RF1 AG063903 (Kelleher lab), R01 CA195732 and a Leukemia and Lymphoma Society Specialized Center of Excellence Grant (Licht Lab) and R44 GM116584 and R44 CA212733 (EpiCypher). LFS is a Gilliam Fellow of the Howard Hughes Medical Institute. Research in this publication is also supported by Thermo Fisher Scientific and a fellowship associated with the Chemistry of Life Processes Predoctoral Training Grant T32 GM105538 at Northwestern University. We thank SCIEX for their support including Dr. Fang Wang for the valuable discussions and insightful suggestions throughout this research project.

## Abbreviations

(AmAc): ammonium acetate
(BGE): background electrolyte
(CL): conductive liquid
(CZE): capillary zone electrophoresis
(dNuc): designer Nucleosome
(EIE): extracted ion electropherogram
(endoNucs): endogenous nucleosomes
(ESI): electrospray ionization
(FWHM): full-width at half maximum
(HAc): acetic acid
(HCD): higher-energy collisional dissociation
(HCl): hydrochloric acid
(LOD): limit of detection
(ME): methylation equivalents
(MNase): micrococcal nuclease
(MWCO): molecular weight cut-off
(nCZE): native capillary zone electrophoresis
(nMS): native mass spectrometry
(Nuc): mononucleosome
(OT): orbitrap
(PTM): post-translational modification
(QE-EMR): Q Exactive HF MS with Extended Mass Range
(rNuc): recombinant Nucleosome
(SL): separation line
(SNR): signal-to-noise ratio
(TIE): total ion electropherogram
(TDMS): top-down mass spectrometry
(UHMR): Ultra High Mass Range.

