## Supplemental Information 1 for "Separation and Characterization of Endogenous Nucleosomes by Native Capillary Zone Electrophoresis – Top-Down Mass Spectrometry (nCZE-TDMS)"

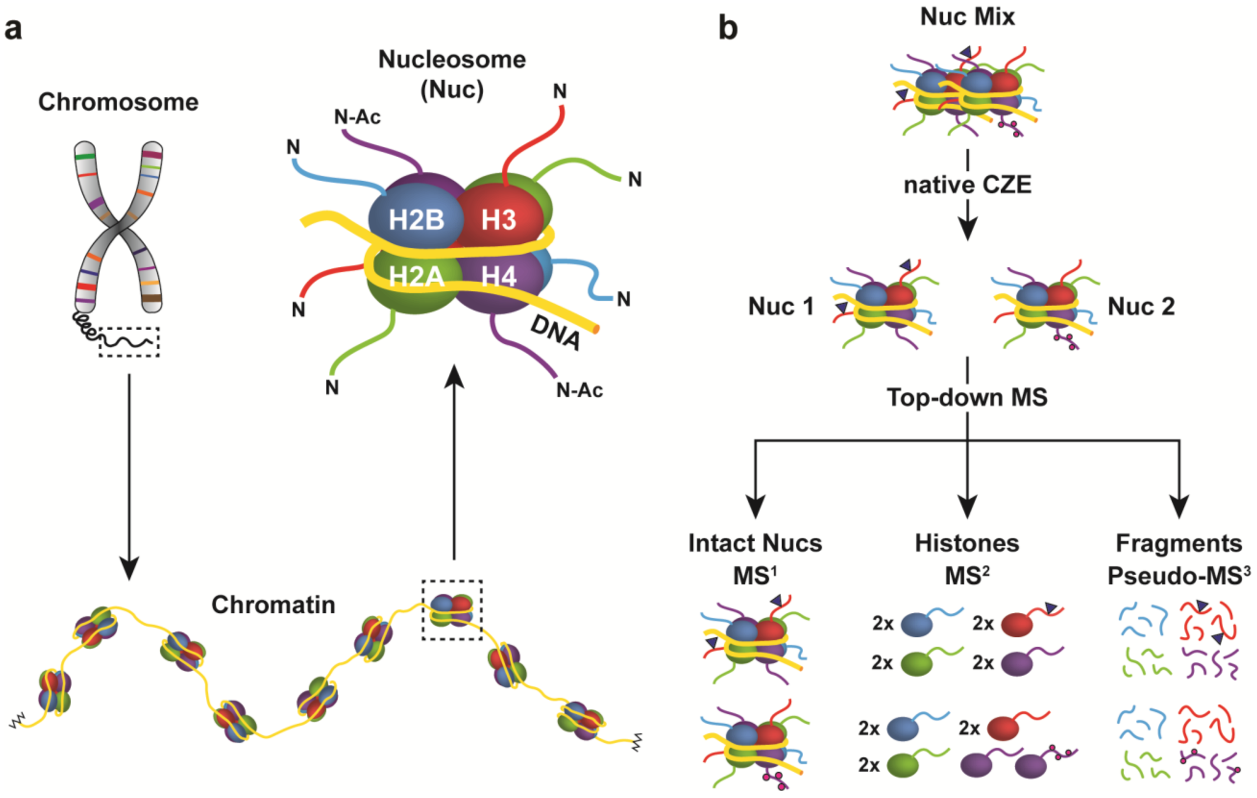


**Figure S1:** (a) Scheme of chromosome, chromatin, and nucleosome (Nuc). Chromosomes are macrostructures based on assemblages of chromatin, a chain repeating DNA : protein biopolymer. The Nuc, the smallest repeating unit, consists of ~147 bp DNA wrapped around two copies each of the core histones: H2A, H2B, H3, and H4. (b) Workflow of nCZE-TDMS system: Separation of different intact Nucs (e.g. proteoforms) in the nCZE dimension. TDMS analysis then follows in three steps: intact Nucs ($MS^1$), histone subunits ($MS^2$), and peptide fragments (pseudo-$MS^3$).

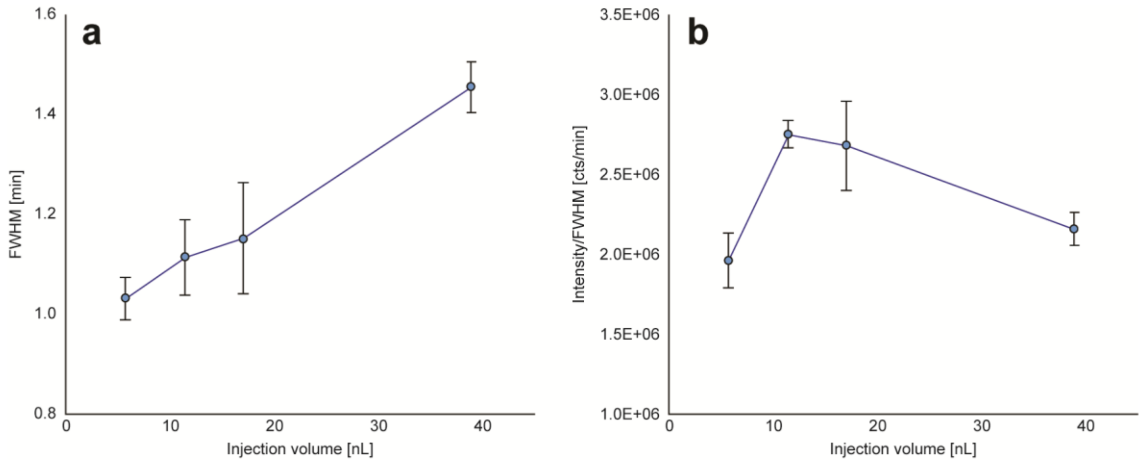


**Figure S2.** Optimization of injection volume. Different volumes of H3G34V (c = 0.75 µM, n = 3), ranging from ~5.7 to 38.0 nL (2.5 psi, 15 to 100 sec) were injected, which corresponds to roughly 0.9 to 6.0% of the total capillary volume. (**a**) Full-width half maximum (FWHM) was plotted against injection volume. (**b**) Ratio between intensity and FWHM was plotted against injection volume.

Optimization of injection volume: As would be expected, CZE peak widths increase proportional with the applied injection volume (**Figure S2a**). One appropriate measure to evaluate peak shape is to look at the ratio between peak intensity and full width half maximum (FWHM). The highest ratio was observed at 11.4 nL injection volume (**Figure S2b**). In addition, there was no significant improvement of absolute intensity observed for higher injection volumes. For this reason, 11.4 nL (2.5 psi, 30 sec) was selected for further experiments.

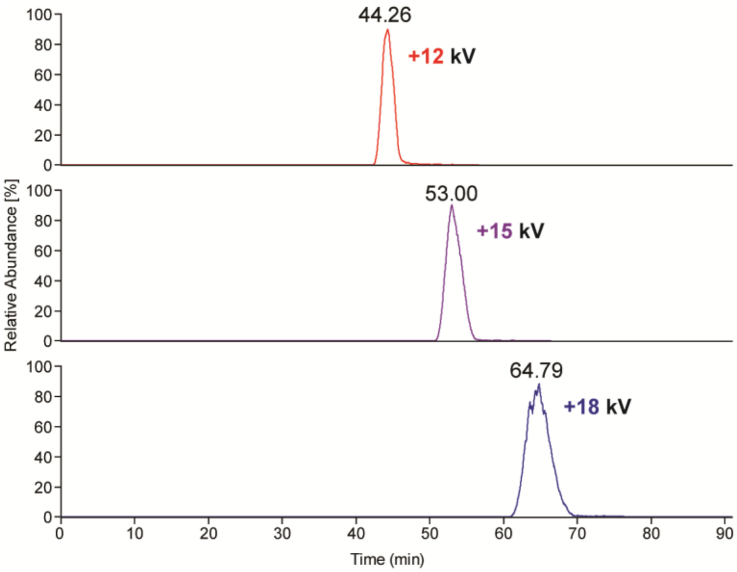


**Figure S3.** Evaluation of influence of separation voltage on nucleosome migration. H2BK120ub (c = 500 nM) was analyzed at three different voltage settings: +12, +15, and +18 kV. The other method parameter settings were kept constant, including 40 mM AmAc (pH ≈ 6.8) as BGE and a supplemental pressure of 2.5 psi. The systematic increase in migration time by changing the HV from +12 to +18 kV, while maintaining other parameters, is a strong indication that net charge of the nucleosome complexes is negative under these conditions.

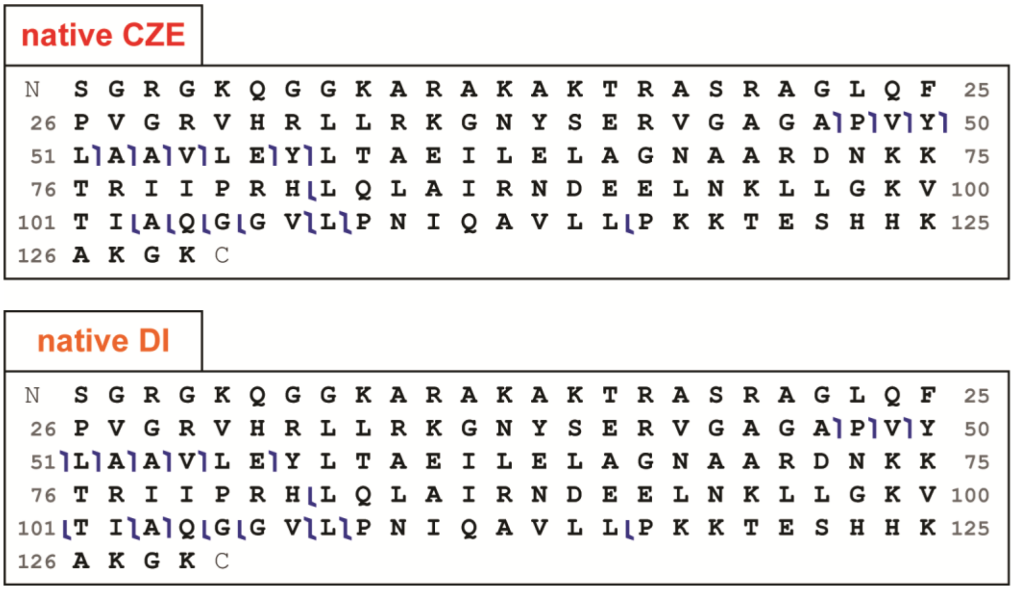


**Figure S4.** Comparison of fragmentation performance between native CZE and native direct infusion (DI) for the analysis of histone H2A after ejection from H3K27me3 nucleosomes by pseudo $MS^3$ TD experiments. The same number of mass spectra were averaged for each technique for comparison. Fragmentation profiles and coverage were determined to be similar between these two approaches and most fragments were picked up by both techniques. This indicates no significant loss of fragmentation quality due to the front end CZE separation.

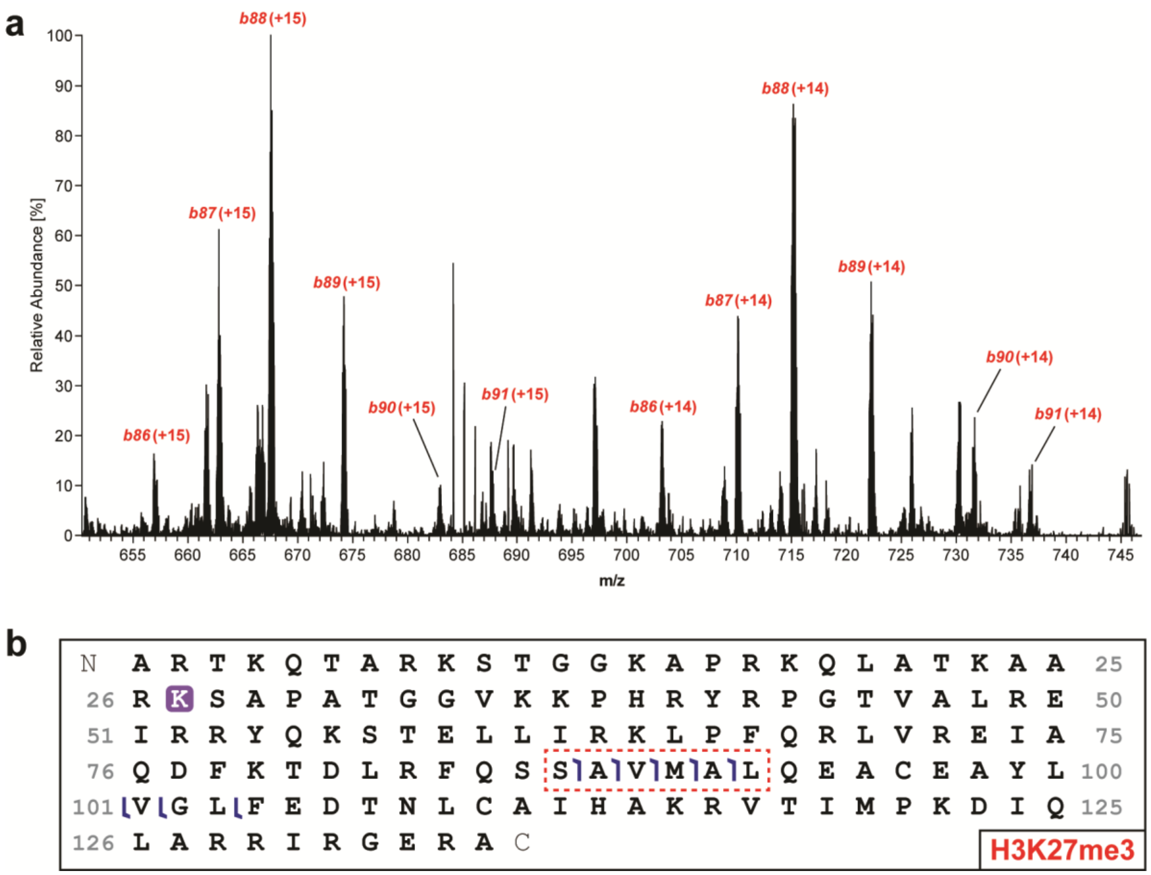


**Figure S5.** (**a**) Highlighted section of pseudo-MS$^3$ mass spectrum showing diagnostic trimethylated *b*-ions after ejection and fragmentation (HCD) of histone H3K27me3 from intact Nuc (c = 1 µM). Predominantly, +14 and +15 charge states were observed for these fragment ions. (**b**) Graphical fragment maps of peptides observed from H3K27me3. Detected *b*-ions exhibiting a mass shift matching a tri-methylation are highlighted (red, dashed box).

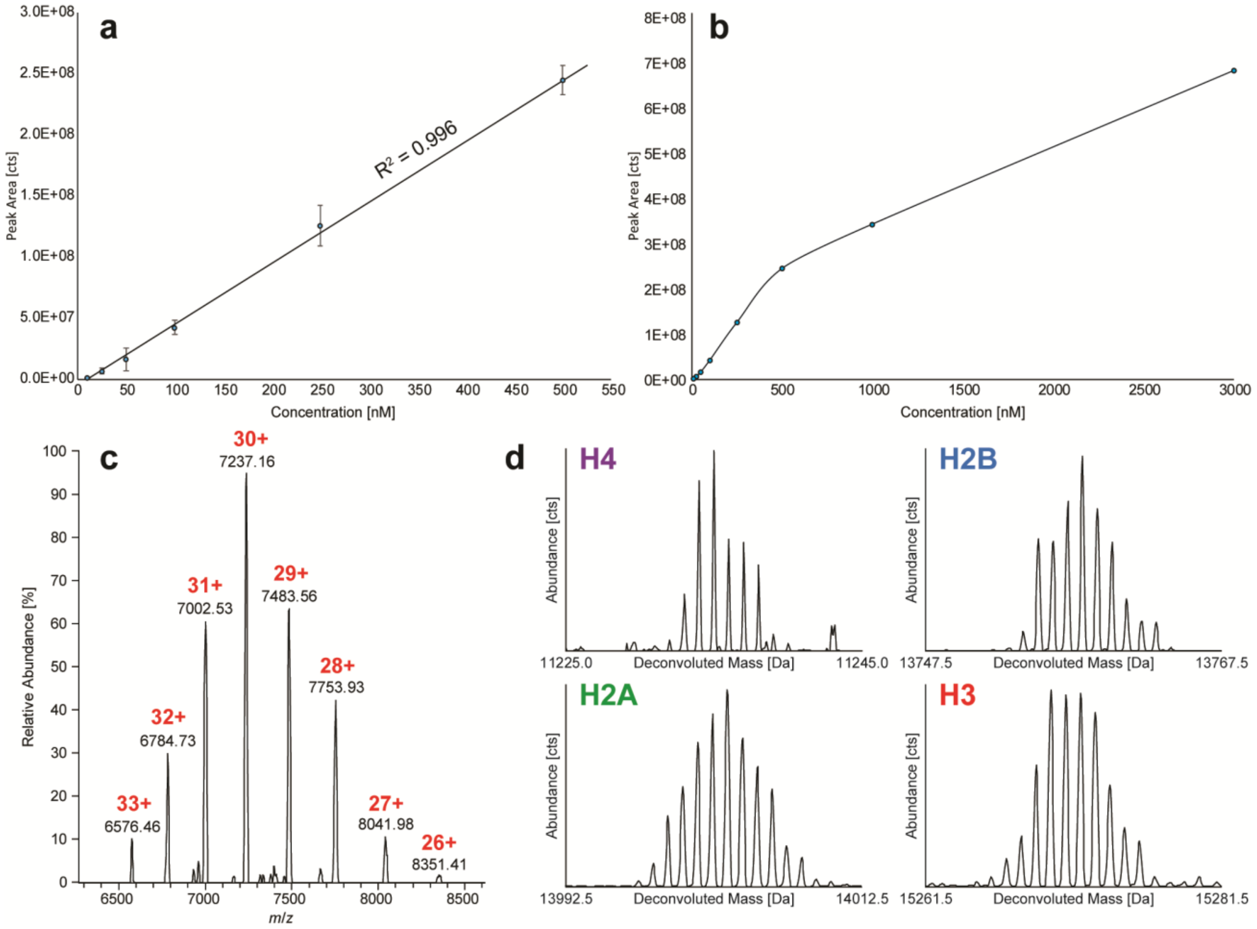


**Figure S6:** Regression and determination of LOD: (a) Linear range was determined as 10 to 500 nM (110 amol to 5.7 fmol) with $R^2$ = 0.996 and (b) dynamic range was observed between 10 to 3000 nM. (c) Mass spectrum of intact nucleosome at lowest analyzed concentration (10 nM) close to LOD with an average S/N of 11.3 ±1.0. (d) Deconvoluted isotopic distributions for histones ejected from unmodified recombinant Nucs measured at 62.5 nM.

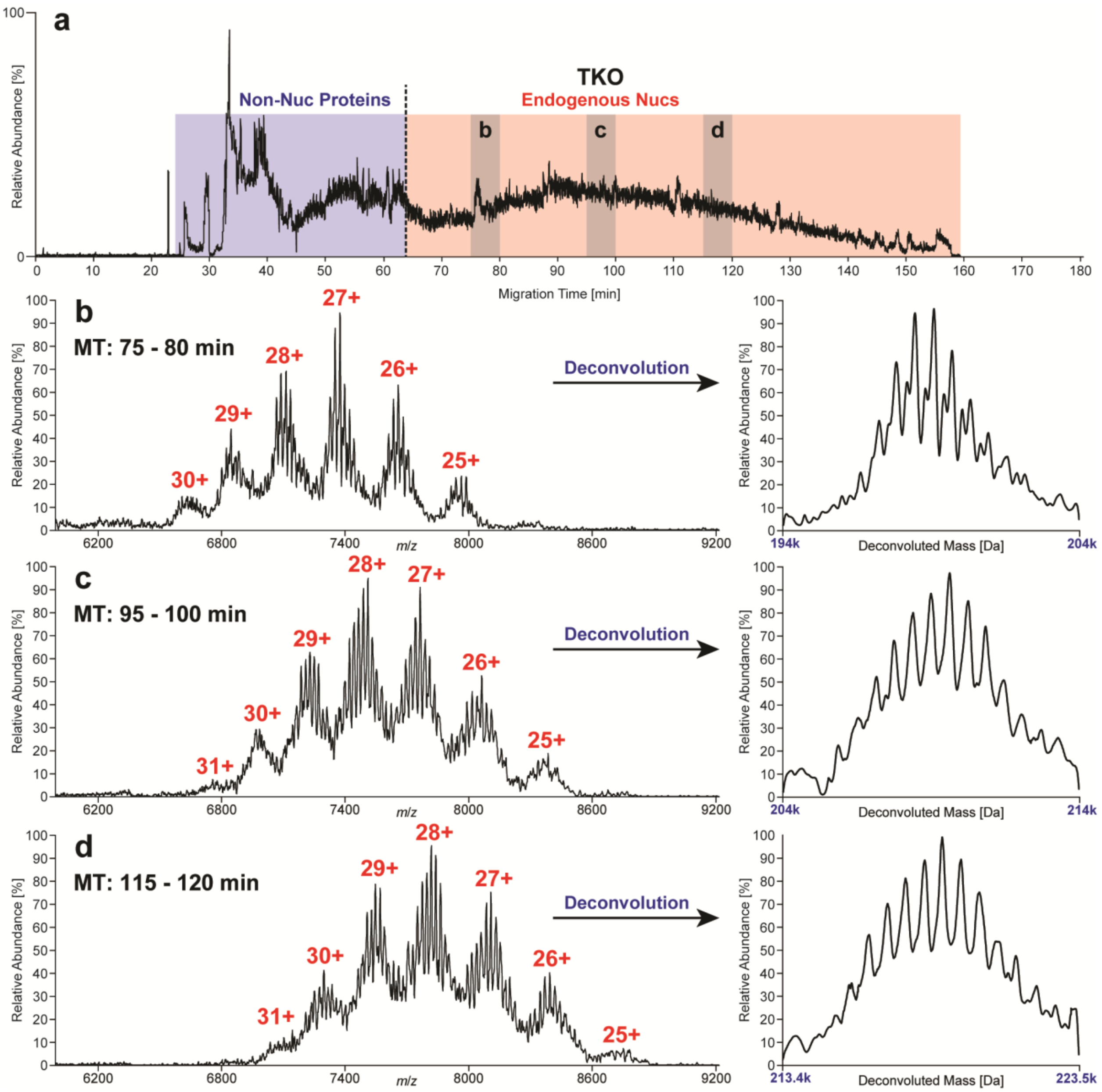


**Figure S7.** Separation of endogenous nucleosomes (endoNucs; ~10 mg/mL) derived from TKO cells. Total Ion Electropherograms (**a**) divided into two major regions: (i) non-Nuc proteins and (ii) endoNucs. Raw and deconvoluted mass spectra (**b-d**) of three exemplary sections (75 – 80 min, 95 – 100 min, 115 -120 min). Several “nucleoforms” were observed in each section. In general, the intact mass of nucleoforms increased with migration time and an average mass shift of 620.2 ±18.5 Da was observed between neighboring peaks, which corresponds to a single base pair difference in length of the associated DNA ($\Delta m_{GC}$ = 618.4 Da, $\Delta m_{AT}$ = 617.4 Da).

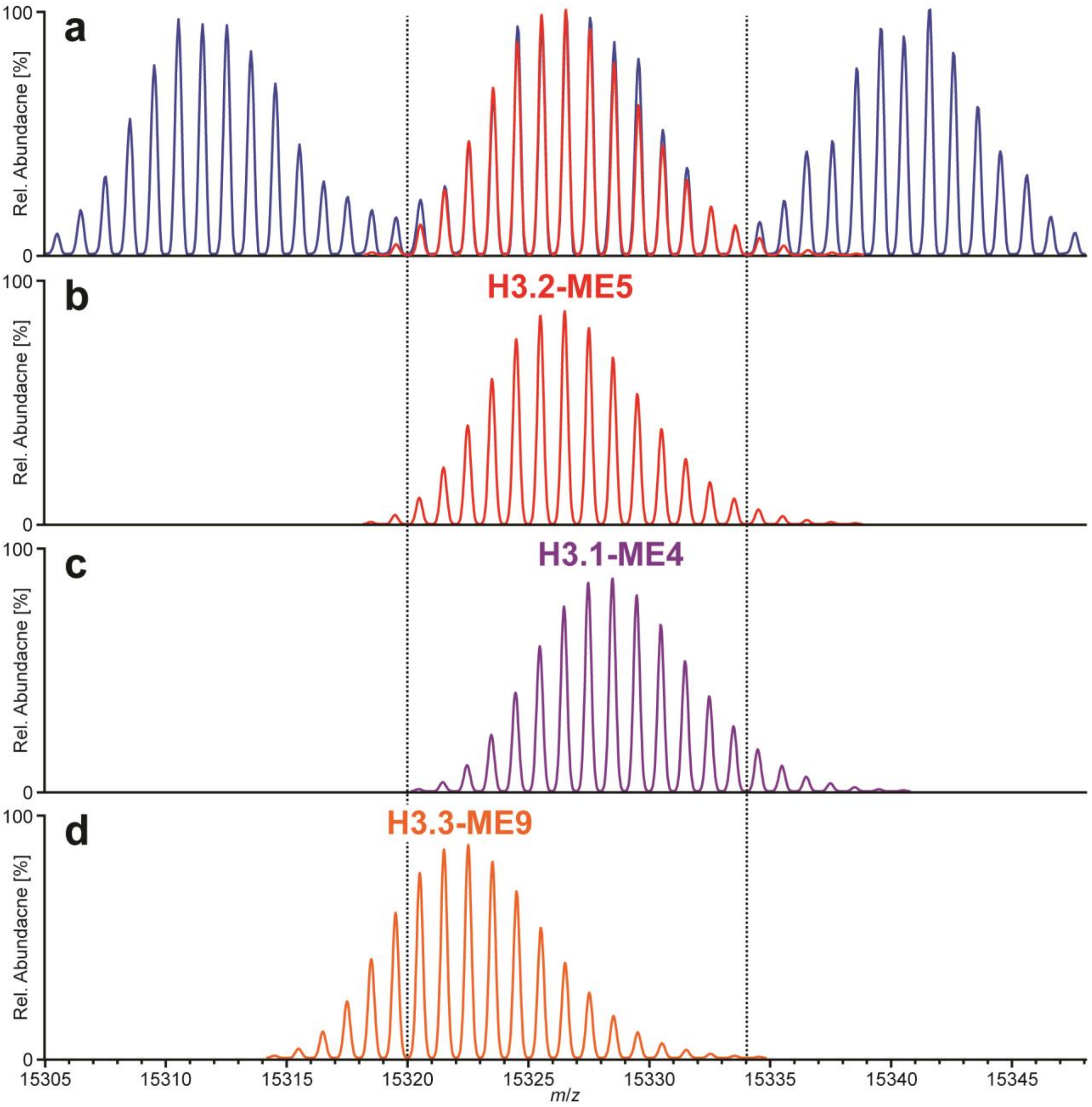


**Figure S8.** Comparison of observed and theoretical isotopic distributions of H3 proteoforms. (**a**) Deconvoluted mass spectrum (blue) of NTKO in the range from 15,305 to 15,350 Da showing the three major H3 proteoforms. The theoretical isotopic distribution of the H3.2 proteoform exhibiting five methyl equivalents (ME) is overlayed (red), indicating a high match with the observed data. Calculated theoretical distributions of proteoforms (**b**) H3.2-ME5, (**c**) H3.1-ME4, and (**d**) H3.3-ME9.

**Table S1:** Overview of steps of the three developed native CE methods. Rinsing steps were performed at 100 psi.

| Step | Capillary | Standard | High-resolution | High-throughput |
|---|---|---|---|---|
| Rinse: 0.1 M HCl | SL | 5 min | 5 min | 3 min |
| Rinse: BGE | CL | 3 min | 3 min | 2 min |
| Rinse: BGE | SL | 5 min | 5 min | 3 min |
| Injection | SL | 2.5 psi, 30 sec | 2.5 psi, 30 sec | 2.5 psi, 30 sec |
| Ramp Up | SL | 5 psi, 1 min | 2.2 psi, 1 min | 15 psi, 1 min |
| Separation | SL | +18 kV, 5 psi, 30 min | +18 kV<br>Synthetic Nucs: 2.2 psi, 115 min<br>EndoNucs: 3.0 psi, 180 min | +18 kV, 15 psi, 6 min |
| Ramp Down | SL | 5 psi, 5 min | 5 psi, 5 min | 15 psi, 4 min |
| Total Run Time | - | ~50 min | ~135 min | ~20 min |

SL = separation line, CL = conductive line

**Table S2:** Parameters of MS methods applied for Nuc characterization using CZE-TDMS.

| **OT Instrument** | **QE-EMR** | | **UHMR** |
|---|---|---|---|
| Mode | $MS^1$ | $MS^2$ | Pseudo-$MS^3$ |
| Scan Range [*m*/*z*] | 500 – 10,000 | 500 – 10,000 | 500 – 12,000 |
| Isolation Range [*m*/*z*] | 2,000 – 10,000 | 6,000 – 9,000 | 1002.5 ± 15 (H2A) |
| Resolution | 7,500; 15,000 | 120,000 | 70,000 |
| In-Source CID [eV] | none | 25 - 50 | - |
| In-Source Trapping | - | - | off |
| Trapping Voltage | - | - | 60 |
| Microscans | 20 | 10 | 10 |
| Max Injection [µs] | 30 | 500 | 1000 |
| AGC target | 3e6 | 3e6 | 1e6 |
| Extended Trapping [eV] | 120 | 120 | - |
| HCD Fragmentation | -10 | -10 | 35 |
| Pressure Regulator | 4 (~1.2e-9 mbar) | 1 (~6e-10 mbar) | - |

**Table S3:** Determination of intra- and inter-day reproducibility for nCZE-TDMS method for nucleosome characterization. Designer nuc H3K27me3 (c = 1 µM) was analyzed five consecutive times on two different days. Relative standard deviation (RSD) values were calculated for migration time (MT), peak area and peak intensity. Peak area and intensity were based on the extracted ion electropherograms (EIEs) of the five most intense charge states.

| **Day 1 (intra-day)** | | | | **Day 2 (intra-day)** | | | |
|---|---|---|---|---|---|---|---|
| **Run** | **MT [min]** | **Area [cts]** | **Intensity [cts]** | **Run** | **MT [min]** | **Area [cts]** | **Intensity [cts]** |
| 1 | 20.44 | $2.60 \cdot 10^{8}$ | $3.80 \cdot 10^{6}$ | 6 | 20.23 | $2.92 \cdot 10^{8}$ | $4.16 \cdot 10^{6}$ |
| 2 | 20.59 | $2.74 \cdot 10^{8}$ | $4.18 \cdot 10^{6}$ | 7 | 20.35 | $2.59 \cdot 10^{8}$ | $3.74 \cdot 10^{6}$ |
| 3 | 20.59 | $2.83 \cdot 10^{8}$ | $4.29 \cdot 10^{6}$ | 8 | 20.20 | $2.53 \cdot 10^{8}$ | $3.82 \cdot 10^{6}$ |
| 4 | 20.42 | $2.63 \cdot 10^{8}$ | $3.78 \cdot 10^{6}$ | 9 | 20.49 | $2.57 \cdot 10^{8}$ | $3.93 \cdot 10^{6}$ |
| 5 | 20.61 | $2.13 \cdot 10^{8}$ | $3.73 \cdot 10^{6}$ | 10 | 20.51 | $2.62 \cdot 10^{8}$ | $3.91 \cdot 10^{6}$ |
| Mean | 20.53 | $2.58 \cdot 10^{8}$ | $3.96 \cdot 10^{6}$ | Mean | 20.36 | $2.62 \cdot 10^{8}$ | $3.91 \cdot 10^{6}$ |
| Stdev | 0.09 | $2.70 \cdot 10^{7}$ | $2.58 \cdot 10^{5}$ | Stdev | 0.14 | $1.71 \cdot 10^{7}$ | $1.58 \cdot 10^{5}$ |
| RSD | 0.45% | 10.46% | 6.53% | RSD | **0.70%** | **6.51%** | **4.04%** |
| | | | | **Day 1 & 2 (inter-day)** | | | |
| | | | | Mean | 20.44 | $2.60 \cdot 10^{8}$ | $3.94 \cdot 10^{6}$ |
| | | | | Stdev | 0.15 | $2.14 \cdot 10^{7}$ | $2.03 \cdot 10^{5}$ |
| | | | | RSD | **0.71%** | **8.22%** | **5.16%** |

**Table S4:** Relative quantification of NTKO/TKO histone proteoforms using nCZE-TDMS. Comparison between histone proteoforms from NTKO (IGH-NSD2; H3K36me2$^{hi}$) and TKO (NSD2$^{+/-}$; H3K36me2$^{lo}$) cells using spectra averaged over the entire endoNuc migration range (n = 3). Statistically significant increases (red) and decreases (blue) of histone proteoforms in TKO samples are highlighted.

| Histone Proteoform | NTKO | | TKO | | t-test | |
|---|---|---|---|---|---|---|
| | Mean | StdDev | Mean | StdDev | p-value | α-value |
| **H4-K20met1** | **3.54%** | **0.02%** | **3.26%** | **0.08%** | **3.91E-03** | 6.25E-03 |
| H4-K20met2 | 3.50% | 0.06% | 3.42% | 0.10% | 0.328 | |
| **H4-NtAc** | **4.22%** | **0.01%** | **5.55%** | **0.17%** | **5.50E-03** | |
| H4-K20met1-NtAc | 8.47% | 0.16% | 8.36% | 0.21% | 0.523 | |
| **H4-K20met2-NtAc** | **50.62%** | **0.63%** | **54.15%** | **0.90%** | **5.19E-03** | |
| H4-K20me3-NtAc / H4-K16Ac-NtAc | 4.23% | 0.16% | 4.38% | 0.12% | 0.273 | |
| **H4-R3met1-K20met3-NtAc / H4 -K20met1-K16Ac-NtAc** | **3.61%** | **0.15%** | **2.93%** | **0.10%** | **2.60E-03** | |
| **H4-K20met2-K16Ac-NtAc** | **21.82%** | **0.18%** | **17.95%** | **0.83%** | **1.42E-03** | |
| H2B PF1 | 2.13% | 0.03% | 1.98% | 0.18% | 0.296 | 6.25E-03 |
| H2B PF2 | 2.73% | 0.06% | 2.52% | 0.18% | 0.123 | |
| H2B PF3 | 5.07% | 0.15% | 4.79% | 0.27% | 0.185 | |
| **H2B.1-K** | **18.30%** | **0.13%** | **20.15%** | **0.12%** | **5.55E-05** | |
| **H2B.1-C/E/F/G/I/O** | **29.95%** | **0.12%** | **31.71%** | **0.44%** | **2.61E-03** | |
| H2B.2-E/F | 34.46% | 0.26% | 32.58% | 0.66% | 0.0102 | |
| **H2B.1-D** | **6.50%** | **0.15%** | **5.44%** | **0.34%** | **7.92E-03** | |
| H2B.1-M | 0.85% | 0.04% | 0.83% | 0.07% | 0.670 | |
| H2A.Z | 1.89% | 0.06% | 2.24% | 0.31% | 0.181 | 3.57E-03 |
| **H2A.1-H-NtAc** | **1.99%** | **0.04%** | **1.53%** | **0.03%** | **6.02E-05** | |
| H2A PF3 | 1.35% | 0.03% | 1.20% | 0.08% | 0.0325 | |
| H2A.2-C-NtAc | 10.57% | 0.02% | 10.72% | 0.21% | 0.350 | |
| H2A.2-B-NtAc | 9.08% | 0.04% | 9.87% | 0.33% | 0.0528 | |
| H2A.J-NtAc | 0.80% | 0.02% | 0.80% | 0.03% | 0.783 | |
| H2A. PF7 | 1.64% | 0.04% | 1.57% | 0.03% | 0.0806 | |
| H2A.1 | 2.99% | 0.05% | 2.82% | 0.07% | 0.0293 | |
| H2A.1C / H2A.1-Q104met1 | 1.38% | 0.02% | 1.27% | 0.04% | 0.0118 | |
| H2A.1-C-Q104met1 | 7.14% | 0.28% | 6.57% | 0.21% | 0.0469 | |
| H2A.1-B/E | 50.14% | 0.55% | 52.62% | 1.32% | 0.0394 | |
| **H2A.1-C-NtAc** | **6.96%** | **0.22%** | **5.94%** | **0.11%** | **1.91E-03** | |
| **H2A.1-B/E-Q104met1** | **2.21%** | **0.07%** | **1.61%** | **0.09%** | **7.23E-04** | |
| **H2A.1-B/E-NtAc** | **1.84%** | **0.01%** | **1.24%** | **0.06%** | **2.66E-03** | |
| H3.3-ME3 | 1.64% | 0.13% | 2.13% | 0.10% | 6.61E-03 | 5.00E-03 |
| **H3.3-ME4** | **3.11%** | **0.19%** | **3.88%** | **0.07%** | **2.66E-03** | |
| **H3.3-ME5** | **4.33%** | **0.10%** | **5.29%** | **0.05%** | **1.26E-04** | |
| **H3.2-ME2** | **5.89%** | **0.18%** | **7.75%** | **0.10%** | **8.74E-05** | |
| **H3.2-ME3** | **11.22%** | **0.25%** | **15.21%** | **0.39%** | **1.20E-04** | |
| **H3.2-ME4** | **20.17%** | **0.14%** | **28.56%** | **0.35%** | **2.64E-06** | |
| H3.2-ME5 | 21.94% | 0.12% | 21.22% | 0.26% | 1.25E-02 | |
| **H3.2-ME6** | **19.71%** | **0.17%** | **9.92%** | **0.34%** | **1.52E-06** | |
| **H3.2-ME7** | **9.30%** | **0.20%** | **4.47%** | **0.19%** | **7.18E-06** | |
| **H3.2-ME8** | **2.69%** | **0.28%** | **1.57%** | **0.08%** | **2.55E-03** | |

PF = proteoform, met = methylation, Ac = acetylation, NtAc = N-terminal acetylation, ME = methyl equivalent

**Table S5:** Relative quantification of NTKO/TKO histones using nCZE-TDMS. Comparison between histones from NTKO and TKO cells using spectra averaged over the entire endoNuc migration range (n = 3).

| **Histone** | **NTKO** | **TKO** |
|---|---|---|
| H4 | 6.04 ±0.07% | 5.81 ±0.24% |
| H2B | 47.24 ±0.74% | 47.02 ±2.42% |
| H2A | 36.20 ±0.60% | 36.54 ±2.00% |
| H3 | 10.52 ±0.24% | 10.63 ±0.75% |

**Table S6:** Comparison between front, middle and end section of the endoNuc migration time window for NTKO/TKO endoNuc samples analyzed by nCZE-TDMS. Electropherograms were divided into three equally long sections and deconvoluted.

| **Histone** | **NTKO** | | | **TKO** | | |
|---|---|---|---|---|---|---|
| | Front | Middle | End | Front | Middle | End |
| H4 | 5.97 ±0.07% | 5.78 ±0.08% | 7.38 ±0.21% | 5.93 ±0.31% | 5.52 ±0.16% | 6.48 ±0.44% |
| H2B | 38.16 ±0.95% | 51.68 ±0.53% | 58.66 ±0.85% | 38.07 ±1.67% | 52.06 ±1.29% | 59.01 ±3.83% |
| H2A | 42.42 ±0.67% | 33.49 ±0.51% | 27.25 ±0.81% | 42.42 ±1.97% | 33.31 ±0.97% | 28.66 ±3.04% |
| H3 | 13.45 ±0.29% | 9.05 ±0.16% | 6.70 ±0.22% | 13.58 ±0.62% | 9.10 ±0.47% | 5.85 ±1.00% |

**Table S7:** Comparison between front, middle and end section of the endoNuc migration time window for **NTKO** (IGH-NSD2; H3K36me2$^{hi}$) endoNuc samples analyzed by nCZE-TDMS. t-tests were performed between front and end sections. Statistically significant increases (red) and decreases (blue) of histone proteoforms in TKO samples are highlighted.

| **Histone Proteoform** | **Front Secion** | | **End Section** | | **t-test** | |
|---|---|---|---|---|---|---|
| | Mean | StdDev | Mean | StdDev | p-value | α-value |
| **H4-K20met1** | **4.29%** | **0.16%** | **2.63%** | **0.08%** | **8.43E-05** | |
| **H4-K20met2** | **3.90%** | **0.22%** | **2.86%** | **0.12%** | **2.07E-03** | |
| H4-NtAc | 4.25% | 0.32% | 4.20% | 0.08% | 0.817 | |
| **H4-K20met1-NtAc** | **8.93%** | **0.24%** | **7.26%** | **0.26%** | **1.21E-03** | |
| **H4-K20met2-NtAc** | **49.63%** | **0.65%** | **56.40%** | **1.15%** | **8.77E-04** | 6.25E-03 |
| H4-K20me3-NtAc / H4-K16Ac-NtAc | 4.22% | 0.33% | 3.66% | 0.36% | 0.0797 | |
| **H4-R3met1-K20met3-NtAc / H4 -K20met1-K16Ac-NtAc** | **4.32%** | **0.23%** | **2.50%** | **0.06%** | **3.67E-04** | |
| H4-K20met2-K16Ac-NtAc | 20.64% | 0.39% | 20.48% | 0.28% | 0.596 | |
| H2B PF1 | 2.08% | 0.05% | 1.76% | 0.11% | 0.0110 | |
| H2B PF2 | 2.66% | 0.09% | 2.30% | 0.11% | 0.0126 | |
| **H2B PF3** | **5.38%** | **0.19%** | **4.08%** | **0.22%** | **3.06E-03** | |
| **H2B.1-K** | **18.48%** | **0.13%** | **17.74%** | **0.11%** | **9.04E-04** | |
| **H2B.1-C/E/F/G/I/O** | **29.98%** | **0.14%** | **31.12%** | **0.27%** | **2.95E-03** | 6.25E-03 |
| **H2B.2-E/F** | **34.01%** | **0.30%** | **36.86%** | **0.53%** | **1.28E-03** | |
| H2B.1-D | 6.33% | 0.19% | 5.71% | 0.27% | 0.0313 | |
| **H2B.1-M** | **1.17%** | **0.06%** | **0.42%** | **0.02%** | **2.70E-05** | |
| H2A.Z | 1.66% | 0.08% | 1.36% | 0.09% | 0.0110 | |
| H2A.1-H-NtAc | 1.96% | 0.04% | 1.77% | 0.06% | 7.85E-03 | |
| H2A PF3 | 1.34% | 0.05% | 1.19% | 0.04% | 0.0142 | |
| **H2A.2-C-NtAc** | **10.14%** | **0.04%** | **10.85%** | **0.13%** | **7.55E-04** | |
| H2A.2-B-NtAc | 8.87% | 0.07% | 8.99% | 0.17% | 0.319 | |
| **H2A.J-NtAc** | **0.80%** | **0.03%** | **0.66%** | **0.03%** | **1.40E-03** | |
| **H2A. PF7** | **1.76%** | **0.06%** | **1.19%** | **0.04%** | **1.92E-04** | |
| **H2A.1** | **3.28%** | **0.13%** | **2.18%** | **0.08%** | **2.54E-04** | 3.57E-03 |
| **H2A.1C / H2A.1-Q104met1** | **1.49%** | **0.04%** | **1.06%** | **0.04%** | **1.43E-04** | |
| **H2A.1-C-Q104met1** | **7.65%** | **0.29%** | **5.38%** | **0.22%** | **4.11E-04** | |
| **H2A.1-B/E** | **49.09%** | **0.66%** | **56.73%** | **1.07%** | **4.57E-04** | |
| H2A.1-C-NtAc | 7.12% | 0.25% | 5.93% | 0.33% | 7.41E-03 | |
| **H2A.1-B/E-Q104met1** | **2.35%** | **0.09%** | **1.62%** | **0.11%** | **8.27E-04** | |
| **H2A.1-B/E-NtAc** | **2.42%** | **0.02%** | **1.09%** | **0.04%** | **8.16E-07** | |
| H3.3-ME3 | 1.73% | 0.10% | 1.73% | 0.21% | 0.978 | |
| H3.3-ME4 | 3.25% | 0.14% | 3.11% | 0.14% | 0.284 | |
| H3.3-ME5 | 4.36% | 0.04% | 4.20% | 0.36% | 0.531 | |
| **H3.2-ME2** | **5.90%** | **0.10%** | **5.31%** | **0.14%** | **3.83E-03** | |
| H3.2-ME3 | 11.17% | 0.20% | 11.65% | 0.46% | 0.177 | |
| H3.2-ME4 | 20.17% | 0.04% | 19.50% | 0.20% | 0.0264 | 5.00E-03 |
| **H3.2-ME5** | **21.49%** | **0.10%** | **22.73%** | **0.31%** | **2.61E-03** | |
| H3.2-ME6 | 19.76% | 0.19% | 20.65% | 0.27% | 9.37E-03 | |
| H3.2-ME7 | 9.40% | 0.33% | 8.96% | 0.26% | 0.145 | |
| H3.2-ME8 | 2.76% | 0.25% | 2.17% | 0.35% | 0.0734 | |

PF = proteoform, met = methylation, Ac = acetylation, NtAc = N-terminal acetylation, ME = methyl equivalent

**Table S8:** Comparison between front, middle and end section of the endoNuc migration time window for **TKO** ($NSD2^{+/-}$; $H3K36me2^{lo}$) endoNuc samples analyzed by nCZE-TDMS. t-tests were performed between front and end sections. Significant increases (red) and decreases (blue) of histone proteoforms at higher migration times are highlighted.

| Histone Proteoform | Front Secion | | End Section | | t-test | |
|---|---|---|---|---|---|---|
| | Mean | StdDev | Mean | StdDev | p-value | α-value |
| **H4-K20met1** | **3.63%** | **0.11%** | **2.48%** | **0.07%** | **1.05E-04** | 6.25E-03 |
| **H4-K20met2** | **3.72%** | **0.05%** | **2.86%** | **0.14%** | **5.05E-04** | |
| H4-NtAc | 5.35% | 0.06% | 5.57% | 0.34% | 0.380 | |
| H4-K20met1-NtAc | 8.65% | 0.40% | 7.57% | 0.24% | 0.0164 | |
| H4-K20met2-NtAc | 52.00% | 2.04% | 60.95% | 2.68% | 0.0100 | |
| H4-K20me3-NtAc / H4-K16Ac-NtAc | 4.79% | 0.23% | 3.19% | 0.57% | 0.0109 | |
| H4-R3met1-K20met3-NtAc / H4 -K20met1-K16Ac-NtAc | 3.64% | 0.58% | 1.93% | 0.23% | 8.91E-03 | |
| H4-K20met2-K16Ac-NtAc | 18.22% | 1.39% | 15.45% | 1.89% | 0.109 | |
| H2B PF1 | 1.96% | 0.20% | 1.69% | 0.11% | 0.112 | 6.25E-03 |
| H2B PF2 | 2.48% | 0.22% | 2.06% | 0.20% | 0.0710 | |
| H2B PF3 | 4.87% | 0.30% | 3.80% | 0.40% | 0.0211 | |
| H2B.1-K | 20.10% | 0.26% | 19.82% | 0.22% | 0.231 | |
| H2B.1-C/E/F/G/I/O | 31.69% | 0.55% | 33.12% | 0.40% | 0.0217 | |
| H2B.2-E/F | 32.35% | 0.86% | 34.72% | 0.76% | 0.0231 | |
| H2B.1-D | 5.46% | 0.40% | 4.30% | 0.42% | 0.0255 | |
| **H2B.1-M** | **1.10%** | **0.11%** | **0.48%** | **0.07%** | **1.19E-03** | |
| H2A.Z | 2.14% | 0.33% | 1.67% | 0.17% | 0.0944 | 3.57E-03 |
| H2A.1-H-NtAc | 1.50% | 0.06% | 1.37% | 0.11% | 0.143 | |
| H2A PF3 | 1.31% | 0.09% | 1.00% | 0.05% | 0.0220 | |
| H2A.2-C-NtAc | 10.55% | 0.11% | 10.21% | 0.75% | 0.519 | |
| H2A.2-B-NtAc | 9.71% | 0.36% | 9.47% | 0.51% | 0.552 | |
| H2A.J-NtAc | 0.84% | 0.06% | 0.67% | 0.04% | 0.0132 | |
| H2A. PF7 | 1.71% | 0.01% | 1.35% | 0.16% | 0.0571 | |
| **H2A.1** | **3.06%** | **0.12%** | **2.31%** | **0.17%** | **3.13E-03** | |
| **H2A.1C / H2A.1-Q104met1** | **1.36%** | **0.04%** | **1.00%** | **0.03%** | **3.45E-04** | |
| **H2A.1-C-Q104met1** | **7.00%** | **0.22%** | **4.96%** | **0.13%** | **1.58E-04** | |
| **H2A.1-B/E** | **51.52%** | **1.58%** | **59.37%** | **1.40%** | **2.98E-03** | |
| **H2A.1-C-NtAc** | **6.23%** | **0.17%** | **4.37%** | **0.27%** | **5.66E-04** | |
| H2A.1-B/E-Q104met1 | 1.71% | 0.16% | 1.09% | 0.20% | 0.0131 | |
| H2A.1-B/E-NtAc | 1.46% | 0.11% | 1.15% | 0.58% | 0.459 | |
| H3.3-ME3 | 2.17% | 0.12% | 2.30% | 0.27% | 0.493 | 5.00E-03 |
| H3.3-ME4 | 3.79% | 0.18% | 4.09% | 0.30% | 0.209 | |
| H3.3-ME5 | 5.33% | 0.10% | 5.29% | 0.88% | 0.947 | |
| H3.2-ME2 | 7.82% | 0.12% | 7.55% | 0.29% | 0.203 | |
| H3.2-ME3 | 15.24% | 0.58% | 15.45% | 0.56% | 0.686 | |
| H3.2-ME4 | 28.20% | 0.39% | 28.71% | 0.70% | 0.325 | |
| H3.2-ME5 | 21.18% | 0.31% | 20.99% | 0.20% | 0.441 | |
| H3.2-ME6 | 10.16% | 0.27% | 9.58% | 0.19% | 0.0359 | |
| H3.2-ME7 | 4.54% | 0.21% | 4.29% | 0.27% | 0.261 | |
| H3.2-ME8 | 1.56% | 0.16% | 1.74% | 0.34% | 0.449 | |

PF = proteoforms, met = methylation, Ac = acetylation, NtAc = N-terminal acetylation, ME = methylation equivalent

**Discussion S1: Optimization of "higher-resolution" CZE method**

Though partial separation was achieved using the "standard" CZE method, there is still room for significant improvement. Therefore, several different method conditions were evaluated, including ammonium acetate concentration (20 – 40 mM), supplemental pressure (2.2 – 5.0 psi), and pH (6.0 – 8.0). The concentration of AmAc did not affect the overall separation performance significantly (e.g. $R_{20mM}$ = 0.365 vs. $R_{40mM}$ = 0.366). A change in the pH of the BGE did not lead to better separation performance. However, if the pH is too low (≤6.0), no intact nucleosome signal was observed, possibly due to issues with complex stability. Moreover, appreciable metal adduct formation was observed at pH 8.0, decreasing overall signal quality. Given that the Nucs are counter-migrating in the system due to their negative net charge, balancing the supplemental pressure and mobility of the Nucs represents a key factor to improve separation performance. Although migration times and peak widths increase noticeably when decreasing the supplemental pressure from 5.0 psi to 2.2 psi, electrophoretic resolution is significantly improved from R = 0.37 to 1.09 (see **Figure 3**).

**Discussion S2: Optimization of "high-throughput" CZE method**

With potential for widespread deployment of the nCZE-TDMS platform, we developed a "high-throughput" CZE method tailored for the quality control environment during semi-synthetic Nuc manufacturing. For this purpose, we adapted the "standard" CZE method to decrease overall run time but still resolve Nuc-related peaks from salt/matrix compounds (**Figure S9**).

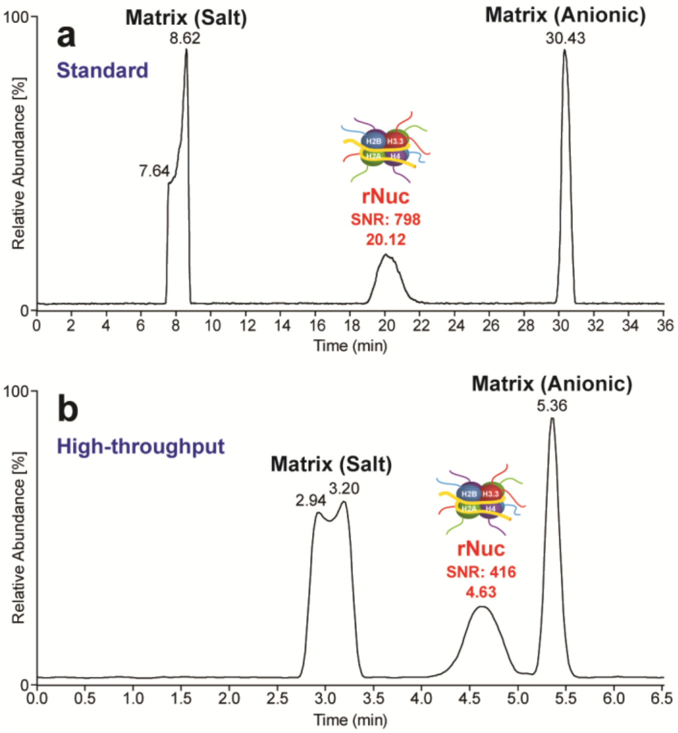


**Figure S9.** Comparison of Total Ion Electropherogram of "standard" (run time: 50 min) and "high-throughput" (run time: 20 min) CZE method analyzing unmodified recombinant Nuc (c = 1 µM). SNR values were determined at FWHM. While the higher-throughput method can baseline-resolve Nuc from matrix, there is a two-fold drop in sensitivity compared to the standard CZE method.

The influence of supplemental pressure was tested in a range between 5.0 and 30 psi and either maintained constant over the course of analysis or varied in different sections of the method. A constant pressure above 15 psi lead to issues regarding resolution of Nuc and matrix compounds.

In addition, varying the pressure during analysis resulted in electrospray process instabilities and thus, potential negative effects on reproducibility. We determined that a constant supplemental pressure of 15 psi, in combination with some changes to the preconditioning steps, yielded stable and reproducible signal, and reduced total run time from 50 to < 20 min per analysis (**Figure S9**). The Nuc peak could be detected in < 5 min using the “high-throughput” CZE, which represents a major improvement versus ~20 min using the “standard” CZE method.
